# Ribociclib as a Potential Multi-Target Inhibitor of Pro-Inflammatory Cytokines: An In Silico Investigation

**DOI:** 10.64898/2026.02.04.703909

**Authors:** Rabita Rahman Era, Akid Ornob

## Abstract

Ribociclib, a selective cyclin-dependent kinase (CDK) 4/6 inhibitor, is approved as a first-line therapy for HR-positive/HER2-negative advanced breast cancer. Emerging evidence suggests that Ribociclib may exert immunomodulatory effects. However, its role in cytokine regulation remains largely unexplored. This study presents a comprehensive in silico investigation of Ribociclib’s interactions with eight key pro-inflammatory cytokines—IL-6, TNF-α, IL-17A, IL-17F, IL-17A/F, IL-1β, MCP-1, and IFN-γ. Computational assessments included molecular docking, molecular dynamics (MD) simulations, MM-GBSA binding free energy calculations, principal component analysis (PCA), and dynamic cross-correlation matrix (DCCM) analyses. Molecular docking and MD simulations indicated strong and stable complex formation with TNF-α, IL-6, MCP-1, IL-1β, and IL-17A/F. MM-GBSA results further showed that Ribociclib formed the most stable complexes with IL-17A/F (ΔG_bind_ = −25.94 kcal/mol) and MCP-1 (ΔG_bind_ = −25.88 kcal/mol), comparable to binding with the CDK-6 (ΔG_bind_ = −36.23 kcal/mol) control protein. PCA and DCCM analyses further supported the stabilizing influence of Ribociclib on these cytokine conformations. Moderate interactions were observed with TNF-α, IL-6, and IFN-γ. Collectively, these findings suggest that Ribociclib may function as a multi-target inhibitor capable of modulating diverse inflammatory pathways, providing a computational foundation for its repurposing as a cost-effective anti-inflammatory therapeutic candidate.

## 1. Introduction

Ribociclib, a cyclin-dependent kinase (CDK) inhibitor, suppresses or averts cancer cell proliferation by targeting CDK4 and CDK6, which are found in proliferating cells such as hematopoietic cells, epithelial cells, fibroblasts, neural progenitor cells, and endothelial cells (1). It is generally used to treat advanced or metastatic hormone receptor-positive, HER2-negative breast cancer in postmenopausal women, frequently in conjunction with hormone treatment such as aromatase inhibitors or selective estrogen receptor modulators (SERMs) (2). The small molecule’s pyridopyrimidine core structure enables it to bind to CDK4/6’s ATP-binding pocket, thereby reducing its activity, and preventing the phosphorylation of the retinoblastoma protein (Rb), causing cell cycle arrest in the G1 phase (3). This efficiently inhibits cell cycle progression and results in decreased tumor development as well as potential cancer cell death (4).

Despite Ribociclib’s recent success as an FDA approved anti-cancer drug, its potential interactions with other molecules remain largely unknown. This lack of thorough understanding not only restricts its broader applicability but also adds to the drug’s high cost. For example, Kisqali, a commercial version of Ribociclib, costs around $7,459 for a packet of 21 tablets at a daily dose of 200 mg (5). Recent studies have investigated Ribociclib’s potential role in modulating immunological pathways (3). Coupling Ribociclib with a TLR4 agonist has been shown to stimulate interleukin-1β (IL-1β) release via an NLRP3- and caspase-dependent mechanism (6). This was shown by combining Ribociclib with a TLR4 agonist in liposomes, which served as an adjuvant in an ovalbumin vaccination model and promoted antigen-specific immunity through interleukin-1 (IL-1β) receptor dependency (6). Furthermore, Chen et al. explored Ribociclib’s potential for treating acute respiratory distress syndrome (ARDS) and neutrophilic inflammation (7). Ribociclib was found to be a novel inhibitor of Phosphodiesterase-4 (PDE-4), which stimulated the cyclic adenosine monophosphate-protein kinase A (cAMP-PKA) pathway and suppressed pro-inflammatory cytokine IL-1β (8). Both preventive and post-treatment with Ribociclib decreased neutrophil infiltration, lung inflammation, and pulmonary damage in mice with Lipopolysaccharide (LPS)-induced ARDS, underlining the drug’s potential in controlling inflammatory responses (7,8). More recently, our prior research on developing a deep learning based virtual drug screening pipeline against the Tumor necrosis factor alpha (TNF-α) protein identified the natural variant of Ribociclib as a potent anti-inflammatory agent based on binding affinity (9). This evidence points towards Ribociclib’s potential activities against multiple pro-inflammatory cytokines.

Pro-inflammatory cytokines are important signaling molecules that are critical for regulating cell growth and differentiation, inducing inflammation, and influencing immunological responses (10). They are important therapeutic targets because they contribute to the pathophysiology of many inflammatory and autoimmune diseases (11). Several disorders have the simultaneous activation of multiple pro-inflammatory cytokines. Rheumatoid arthritis, psoriasis, nonalcoholic fatty liver disease (NAFLD), diabetes, sepsis, and inflammatory bowel disease (IBD) are all characterized by high levels of TNF-α, IL-6, and IL-1β, which contribute to disease development and symptom aggravation (12,13). TNF-α and IL-6 trigger joint inflammation and deterioration in rheumatoid arthritis, whereas IL-1β exacerbates the inflammatory response (14). Similarly, in psoriasis, TNF-α, type I Interferons (IFNs), and the IL-23/IL-17 axis are involved in the promotion of the inflammatory process, with recent findings further elucidating the complex interplay between these cytokines in the pathogenesis of disease (15). In IBD, cytokines such as IL-12 and IL-23 are the primary drivers of persistent intestinal inflammation (16). Evidence suggests that single inhibitors can interact with several cytokines, giving a broader therapeutic strategy (17). Tocilizumab, which is used for rheumatoid arthritis, targets both IL-6 and IL-1β, whereas Adalimumab reduces TNF-α and may impact IL-6 levels in psoriasis (18) (19). Ustekinumab is used to treat inflammatory bowel disease (IBD) by suppressing both IL-12 and IL-23, which further affects downstream cytokines such as IL-17 and Monocyte chemoattractant protein-1 (MCP-1) (20) (21). Whereas biologic therapies with Tocilizumab, Adalimumab, and Ustekinumab have proven effective in inhibiting multiple cytokines, these large molecules are often expensive and hence less accessible to patients. Small molecules act as suitable alternatives as they are relatively cheaper, synthesized with relative ease, and have the potential for oral administration (22).

Notable small molecule multi-target inhibitors include Ruxolitinib and Tofacitinib, whose activities affect specific pathways within an inflammatory course. Ruxolitinib is a Janus Kinase (JAK) inhibitor approved for the treatment of myelofibrosis and polycythemia vera (23). It has potent inhibition of both JAK1 and JAK2 and, to a lesser degree, the signaling of pro-inflammatory cytokines such as IL-1β, IL-6, and TNF-α (24). Tofacitinib has demonstrated therapeutic potential against rheumatoid arthritis and ulcerative colitis due to its direct inhibition of JAK1 and JAK3. The inhibitor also acts directly against several cytokines including IL-2, IL-4, IL-6, and IL-15, and indirectly against TNF-α and IL-1 (25). Research by Ramsay et al. suggests multi-target inhibitors may have fewer side effects than single-target inhibitors because they can provide a more balanced regulation of inflammatory pathways, thus lowering the likelihood of off-target effects associated with more selective therapies (26). Despite their advantages, multi-target cytokine inhibitors are limited due to the stringent chemistry and high costs and time associated in designing such versatile molecules. Repurposing existing drugs with potential anti-inflammatory activities, such as Ribociclib, may provide a viable and cost-effective strategy for discovering common inhibitory agents against pro-inflammatory cytokines.

The first step in drug repurposing is the thorough in silico investigation of its activity against potential new targets (27). As a result, this work aimed at the comprehensive investigation of Ribociclib’s direct interactions with eight major classes of pro-inflammatory cytokines: interferon-γ (IFN-γ), monocyte chemoattractant protein-1 (MCP-1), interleukin-6 (IL-6), interleukin-17A (IL-17A), interleukin-1β (IL-1β), interleukin-17F (IL-17F), tumor necrosis factor-alpha (TNF-α) and interleukin-17A/F (IL-17A/F). Given that Ribociclib is a recognised inhibitor of CDK-6, we utilised CDK-6 as a control protein for comparison. This systematic investigation into the drug’s possible novel functionalities represents the first work of this kind. This work utilized a range of computational techniques to study Ribociclib’s interactions with these cytokines, including molecular docking, molecular dynamics (MD) simulation, MMGBSA, principal component analysis (PCA), and dynamic cross-correlation matrix. Fig 1 displays the methodologies employed, underlining our exhaustive approach in evaluating Ribociclib’s potential as a multi-target inhibitor. More importantly, this work intends to broaden the therapeutic potential of Ribociclib which will make Ribociclib-based therapies cheaper and considerably improve the management of complicated inflammatory and autoimmune illnesses that necessitate a multifaceted response.

**Fig 1.**
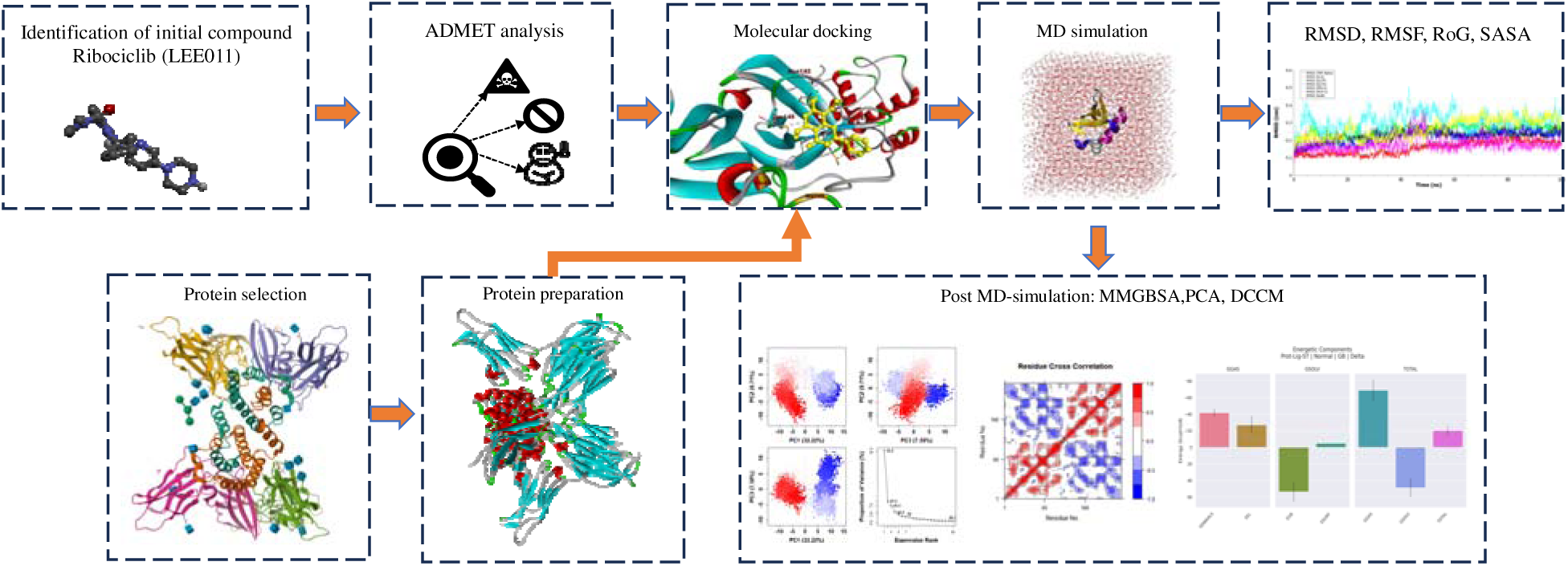
Computational workflow to investigate Ribociclib’s interaction with target proteins: starting with ADMET analysis for drug-like properties, followed by protein preparation and molecular docking for the prediction of binding poses. The MD simulations were carried out to analyze the dynamic behavior and stability of the protein-ligand complexes. Conformational changes were analyzed by PCA and DCCM, while the MMGBSA calculations were performed to gain insight into the binding affinity.

**Fig 2.**
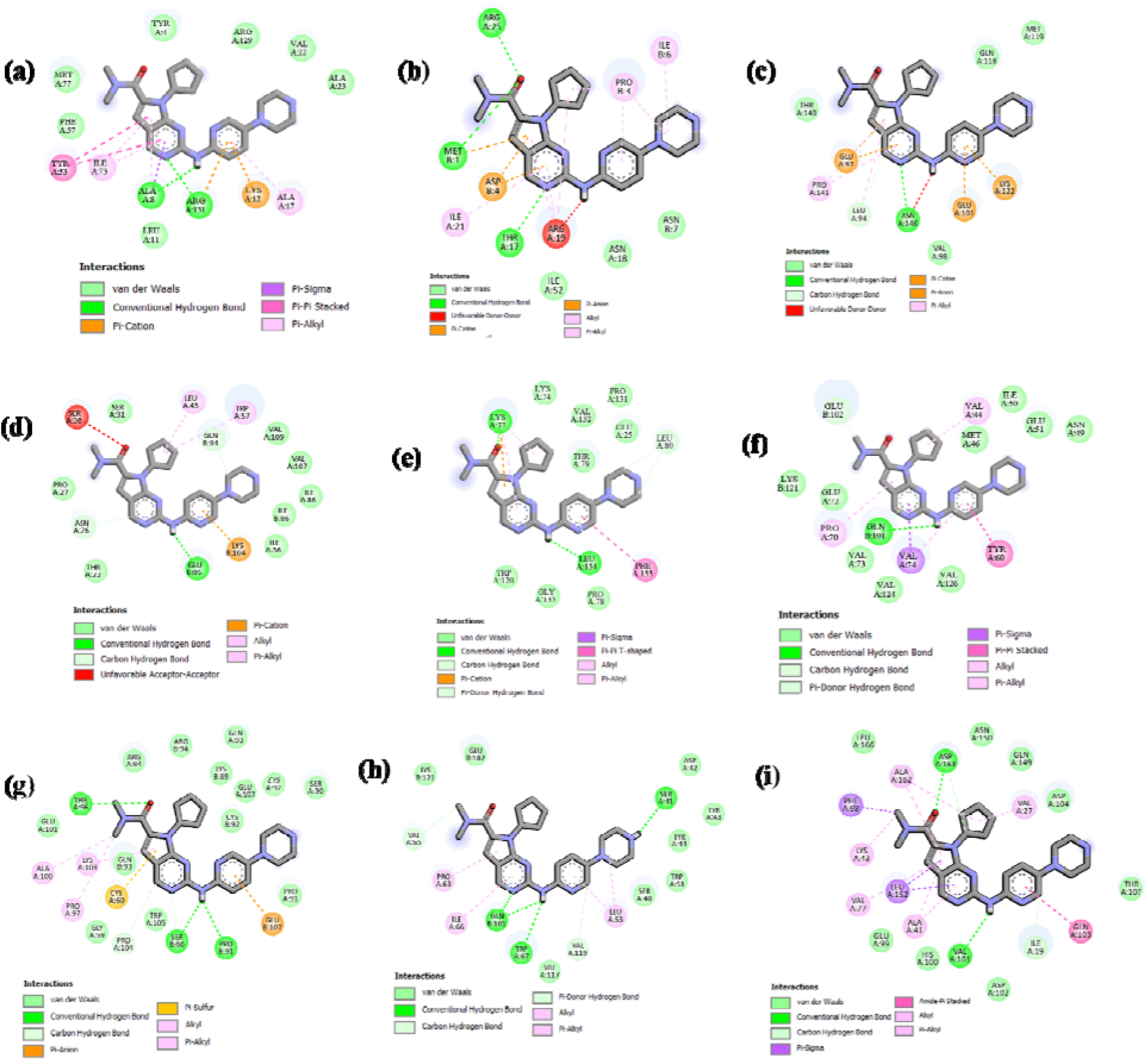
2D Molecular Interactions of Ribociclib with the active sites of (a) IFN-γ (1HIG), (b) MCP-1(1DOK), (c) IL-6 (1ALU), (d) IL-17A (4HR9), (e) IL-1β (9ILB), (f) IL-17F (6HGO), (g) TNF-α (2AZ5), (h) IL-17A/F(5N92). and (i) CDK-6(5L2T).

**Fig 3.**
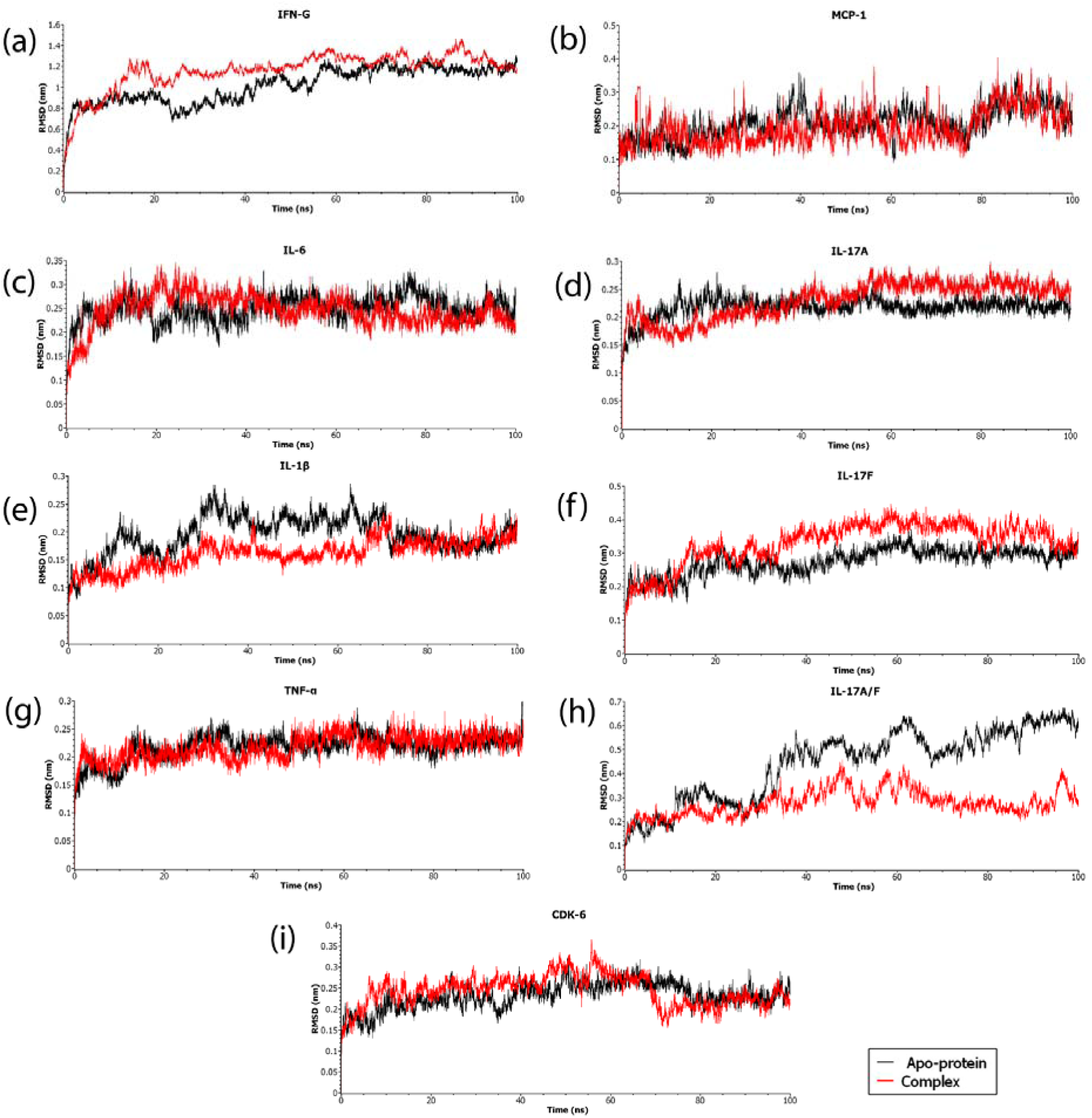
RMSD values plotted for apoprotein (black) and docked complexes(red) (a) IFN-γ (1HIG), (b) MCP-1(1DOK), (c) IL-6 (1ALU), (d) IL-17A (4HR9), (e) IL-1β (9ILB), (f) IL-17F (), (, (g) TNF-α (2AZ5), (h) IL-17A/F(5N92). and (i) CDK-6(5L2T).

## 2. Materials and Methods

### 2.1 Data collection and Preparation

#### 2.1.1 Pro-inflammatory cytokines

IFN-γ, MCP-1, IL-6, IL-17A, IL-1β, IL-17F, and TNF-α represent major classes of pro-inflammatory cytokines. These proteins were chosen to cover a wide range of cytokine signaling pathways, ensuring a comprehensive representation of inflammatory processes. Protein structures were selected based on previous reports of these proteins in structure-activity relationship (SAR), virtual screening, and molecular docking-based studies. The PDB IDs of the selected proteins were 1HIG (IFN-γ), 1DOK (MCP-1), 1ALU (IL-6), 4HR9 (IL-17A), 9ILB (IL-1β), 6HGO (IL-17F), 2AZ5 (TNF-α), 5N92 (IL-17A/F) and 5L2T (CDK-6). The proteins’ FASTA sequences were obtained from the RCSB PDB (28) and uploaded to the Swiss Model server (29) for homology modeling. The best templates were used to generate homology models, which were then downloaded in PDB format. Homology modeling was primarily conducted to address the problem of missing residues. These models were refined in BIOVIA Discovery Studio Visualizer (30) by eliminating water molecules, ligands, and heteroatoms to ensure structural purity. Hydrogen bonds were added to optimize stability and conformation in Chimera (31), followed by energy minimization using Swiss PDB Viewer (32). This was conducted to enhance the protein structures by minimizing steric clashes and unfavorable interactions, thereby improving their overall stability and reliability for molecular docking. The proteins used in this work are summarized in Table 1 and the final prepared proteins are visualized in **Error! Reference source not found.**

**Table 1.**
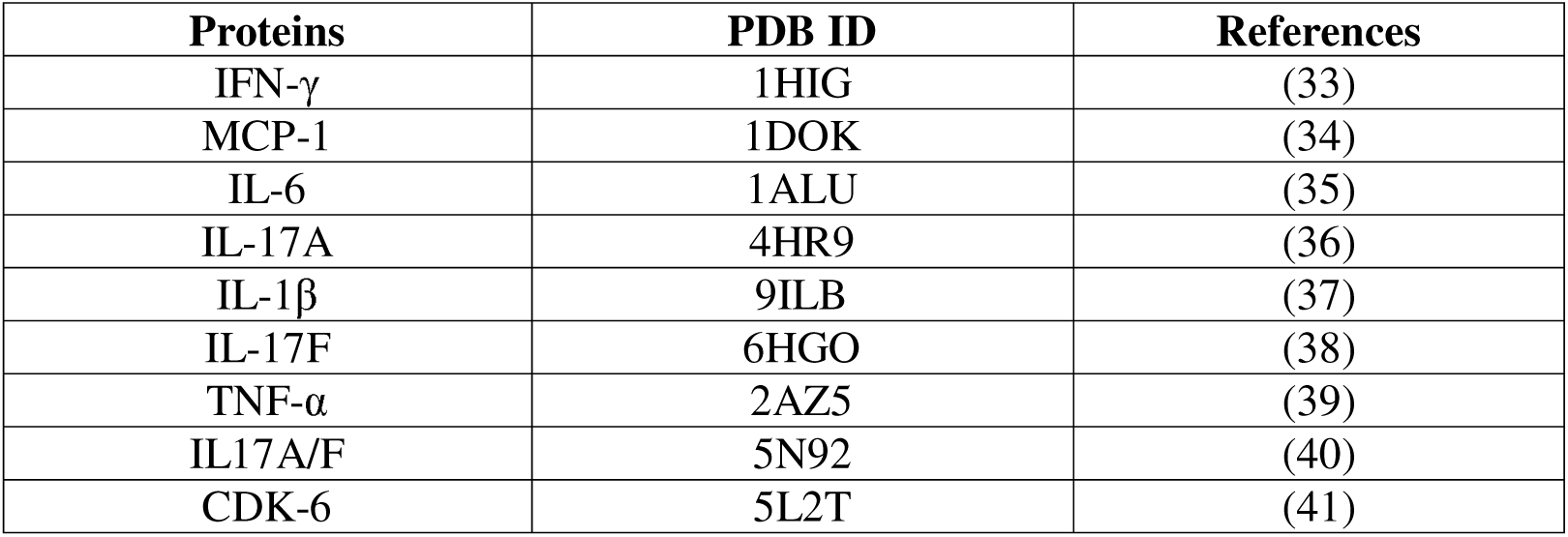
Pro-inflammatory cytokines, their PDB IDs, and corresponding references.

#### 2.1.2 Ligand

Ribociclib, a cyclin-dependent kinase 4/6 (CDK4/6) inhibitor, was chosen as the study’s ligand due to its significance in cancer therapy and possible interactions with pro-inflammatory cytokines as discussed in the previous section. With molecular weight of 434.5 g/mol, the 3D conformer of Ribociclib was obtained from PubChem (42) (PubChem CID: 44631912) in SDF format.

### 2.2 ADMET

ADMET analysis is essential in drug development since it predicts a drug’s effectiveness, safety, and probable interactions with other drugs. Drug candidates’ pharmacokinetic and pharmacodynamic profiles can be enhanced by evaluating these features early in the drug development process, increasing their chances to thrive in clinical trials and regulatory approval. We used two computational tools to evaluate ADMET properties: Protox-II (43) and SwissADME (44). Protox-II is a freely available web-based program with 33 models that provides a broad range of toxicity prediction ability, making it a valuable resource in ADMET analysis. Protox-II was used to evaluate hepatotoxicity, carcinogenicity, immunotoxicity, mutagenicity, cytotoxicity, and LD_50_. SwissADME, a web-based tool, was used to evaluate pharmacokinetic features such as gastrointestinal absorption, blood-brain barrier (BBB) penetration, CYP2C9 inhibition, and drug-likeness. This tool has several unique features, including various input modalities, computation for many molecules at once, and interactive result visualization.

### 2.3 Molecular Docking

The CB-Dock2 tool (45) was used to perform auto-blind docking of proteins with Ribociclib. The processed PDB files of proteins and Ribociclib’s SDF file were submitted to the CB-Dock2 server. For auto-blind docking, CB-Dock2 uses a combination of curvature-based cavity detection and AutoDock Vina-based molecular docking. CB-Dock2 employs the latest version (1.2.0) of AutoDock Vina for template-independent blind docking and utilizes the FitDock method during the docking process. The protein binding pockets were automatically identified, and blind docking was used to explore probable Ribociclib binding locations. CB-Dock2 identified 5-8 potential binding pockets for each protein complex and the poses with the highest affinity were selected.

### 2.4 Molecular Dynamics Simulation

Molecular dynamics (MD) simulation was used to study the stability of protein-ligand complexes at the atomic level. For the molecular dynamics (MD) simulations, GROMACS (version 2023.1) (46) was chosen as the software platform due to its dependability and adaptability in modeling biomolecular systems. The CHARMM36-Jul2022 (47) force field framework was employed, which is well-known for its precision and accurate validation when analyzing molecular interactions within protein-ligand complexes. Simulations were run for 100 nanoseconds (ns) with a time step of 2 femtoseconds, integrating equations of motion over 50,000,000 steps using a leap-frog integrator. Output control methods were used to maintain energy, log files, and compressed coordinate data at 10-picosecond (ps) intervals for complete system monitoring. To ensure stability, bond length constraints involving hydrogen atoms were applied using the Linear Constraint Solver (LINCS) (48) algorithm. Efficient neighbor searching was achieved through the Verlet cutoff scheme (49), with a cutoff distance of 1.2 nanometers (nm) for both electrostatic and van der Waals interactions. Long-range electrostatic interactions were handled using the Particle Mesh Ewald (PME) method (50), with a PME order of 4 and a cutoff distance of 1.2 nm. Na+/Cl-was added to the system to balance the system charge. Temperature was maintained at 300 Kelvin (K) using the V-rescale thermostat, and pressure was kept constant at 1.0 bar using the Parrinello-Rahman Barostat. The system was modeled with three-dimensional periodic boundary conditions (PBC) to mimic an infinite system while maintaining realistic environmental conditions. Dispersion correction was not applied to proteins using the CHARMM36 additive force field. Since simulations continued from a previous NPT equilibration phase, initial velocities were not generated, ensuring a seamless transition into the MD simulation. Using the results of the MD simulation, we conducted several analyses, including the assessment of root-mean-square deviation (RMSD), radius of gyration (Rg), solvent-accessible surface area (SASA), and root-mean-square fluctuation (RMSF) to evaluate the stability and dynamics of the protein-ligand complexes.

### 2.5 MMGBSA

Following the molecular dynamics (MD) simulations, the binding free energy between Ribociclib and the seven target proteins was calculated using the Molecular Mechanics/Generalized Born Surface Area (MM-GBSA) method. For this study, the gmx_mmpbsa tool (51) in GROMACS was used, with last 50 ns trajectory and topology data as input generated from MD simulation. Initially, an index file was created, with system name (“Prot-Lig-ST”), frame range (5000–10,000), and GB model parameters (igb=5, saltcon=0.150). Relevant energy components, including Van der Waals (VDWAALS), Electrostatic (EEL), Generalized Born (EGB), surface (ESURF), gas (GGAS), and solvation (GSOLV) energies, were extracted from each trajectory frame. The collected findings were then analysed using the gmx_mmpbsa ana module to calculate the average, standard deviation, and standard error of the mean for each energy component over all trajectory frames.

### 2.6 Principal component analysis (PCA) and Dynamics cross-correlation matrices (DCCM) analysis

PCA was performed to minimize the dimensionality of the trajectory data and identify the primary movements influencing the proteins’ conformational dynamics. DCCM analysis was employed to evaluate the correlated movements of residues inside proteins. For the final 50 ns of the 100 ns simulation, we used the Bio3D (52) package in RStudio (53) to perform PCA and DCCM analysis. Specifically, we examined the last 5000 frames of the MD simulation trajectory to observe protein dynamics at that stage.

## 3. Results and Discussion

### 3.1 ADMET Analysis

It can be observed from Table 2 that Ribociclib has a high gastrointestinal (GI) absorption rate, indicating effective absorption via the digestive system and raising its bioavailability. Importantly, Ribociclib does not cross the blood-brain barrier (BBB), lowering the likelihood of adverse effects in central nervous system. However, it inhibits CYP2C9, which may affect the metabolism of other drugs and require close monitoring for potential drug interactions. Ribociclib has low scores for hepatotoxicity, carcinogenicity, immunotoxicity, cytotoxicity, and mutagenicity, indicating low risks of liver toxicity, cancer, immune system side effects, cellular toxicity, and mutation induction. Furthermore, the LD_50_ value of 2500 mg/kg indicates Ribociclib’s low acute toxicity. While Ribociclib satisfies Lipinski’s, Egan, Muegge and Veber’s standards for drug likeness, it does not satisfy Ghose’s rule since its molar refractivity exceeds 130. However, since the molar refractivity of Ribociclib is close to 130 (132.5), it has been used clinically and is considered relatively safe according to the toxicity assessment.

**Table 2.**
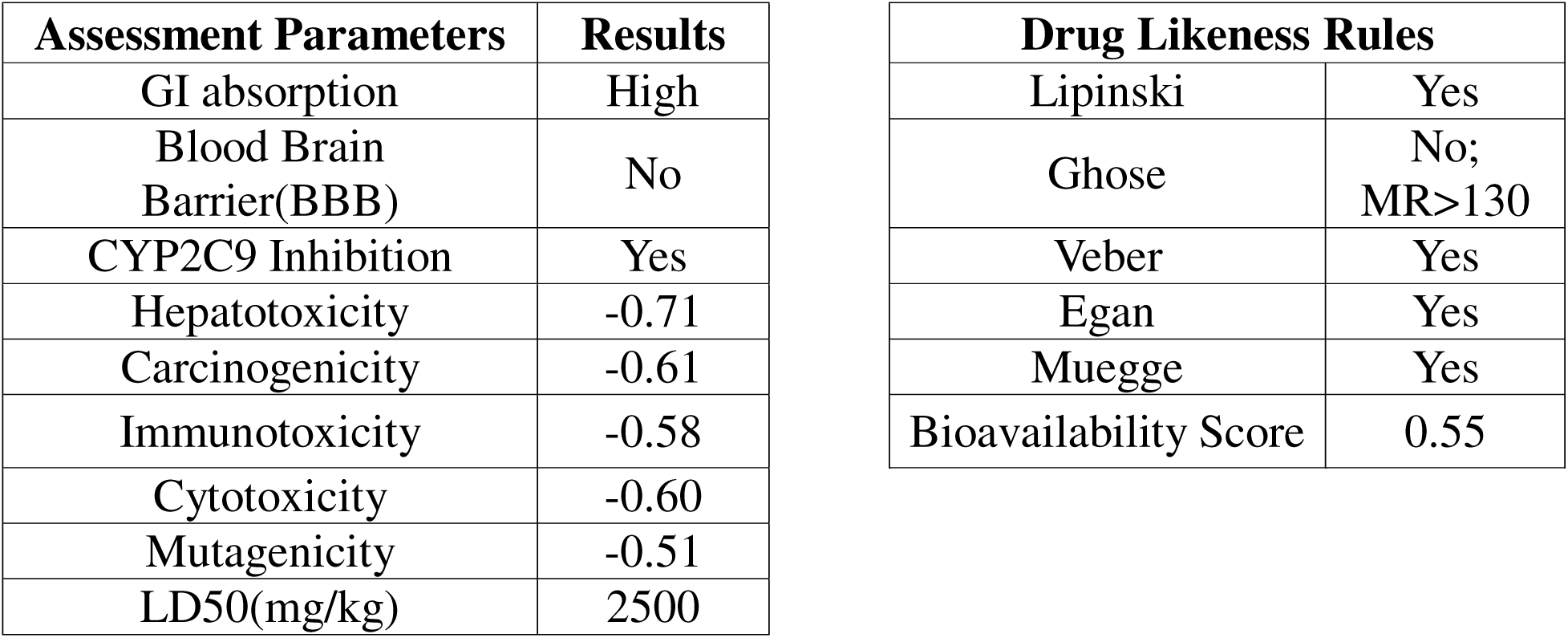
ADMET Analysis of Ribociclib.

### 3.2 Molecular Docking

The molecular docking scores derived from CB-Dock2 revealed Ribociclib’s binding affinity scores with eight target proteins linked in inflammatory processes. Table 3 lists the binding affinity scores in kcal/mol against the protein targets along with CDK-6 control group. Ribociclib has a high affinity for binding for the majority of the target proteins, indicating potential interactions and suggesting effectiveness in inflammatory pathways. TNF-α (2AZ5) exhibited the highest binding affinity (−9.8 kcal/mol), followed by IL-17A/F (5N92) and IL-17F (6HGO), which obtained scores of −8.7 kcal/mol and −8.3 kcal/mol, respectively. Other proteins, including IL-17A (4HR9), IFN-γ (1HIG), IL-1β (9ILB) and MCP-1 (1DOK), demonstrated binding affinities ranging from −8 to −7 kcal/mol. IL-6 (1ALU) had the least favorable binding affinity score with −6.9 kcal.mol. With the CDK-6 (5L2T), Ribociclib showed moderate binding affinity (−7.1kcal/mol). This observation and the specific interactions were confirmed with prior studies (54).

**Table 3.**
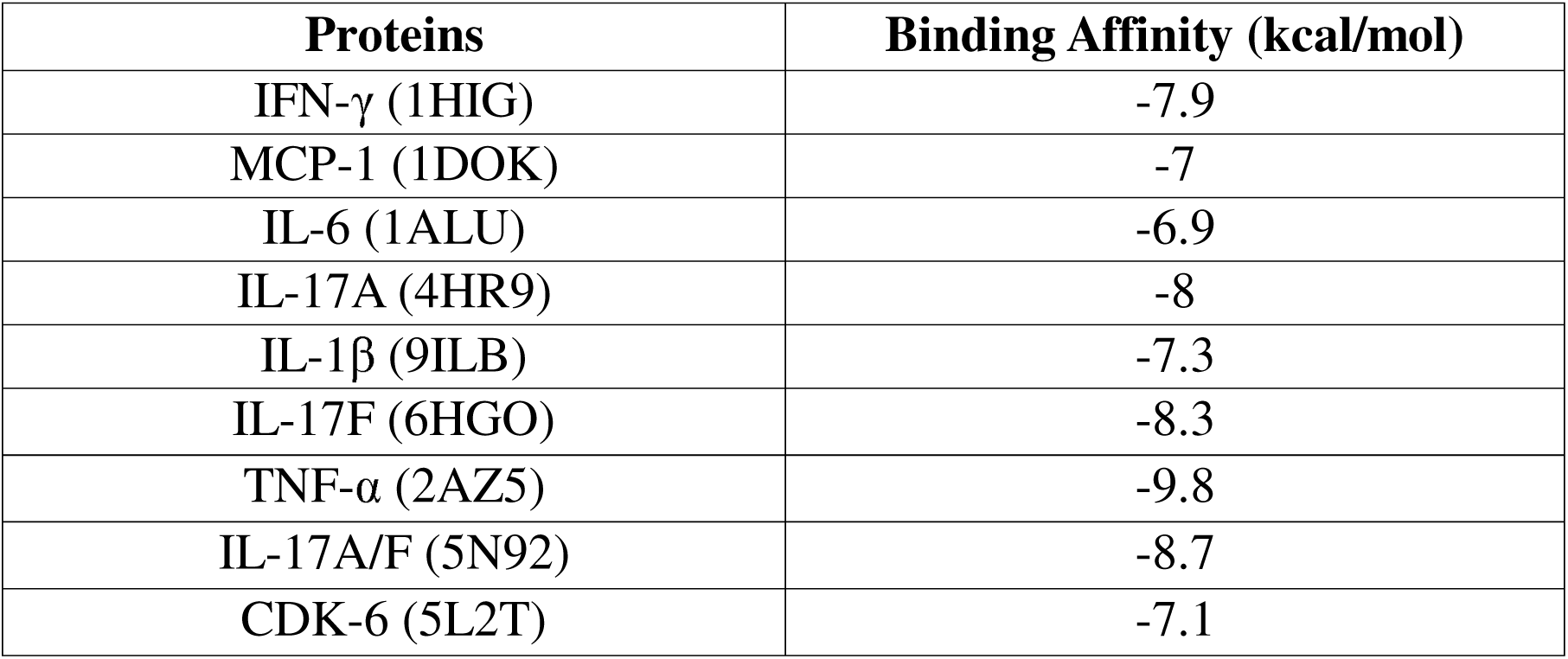
Binding affinity scores from molecular docking.

**Error! Reference source not found.** depicts docked complexes of Ribociclib with eight pro-inflammatory cytokines and CDK-6, providing a visual representation of their binding interactions. Each complex exhibits the spatial arrangement of Ribociclib within the targeted protein’s binding site, revealing potential interaction patterns and binding orientation. The obtained results indicate that Ribociclib interacts in distinct binding pockets within the protein with good conformational stability as can be seen for most proteins. Such intimate positioning indicates that the favorable binding interactions are likely, which further established the suitability of the Ribociclib towards targeting sites within these proteins.

**Error! Reference source not found.** represents the docked complexes’ active sites, with an emphasis on the areas where Ribociclib interacts with the specific amino acids in the target proteins. Overall, we observed the presence of diverse, multiple, and mostly favorable molecular interactions. Ribociclib interacts with the target proteins by a wide range of mechanisms, including van der Waals forces, carbon hydrogen bond, conventional hydrogen bonds, and pi interactions. Van der Waals forces play dominant roles in establishing stability of the docked complexes and underscore the close contact between the Ribociclib and binding sites of the proteins. In addition to forming Van der Waals interactions, TNF-α (2AZ5) exhibited Pi-sulfur bond with CYS60, carbon-hydrogen bond with PRO104, and Pi-anion bond with GLU107. Similarly, in IL-17F (6HGO), the van der Waals forces are supplemented by one conventional hydrogen bonds with GLN101, and pi-sigma interactions with VAL74. The bindings of IL-17A/F (5N92) and IFN-γ (1HIG) are dominated by van der Waals interactions, conventional hydrogen bonds and pi interactions. Ribociclib interacts with MCP-1 (1DOK) by van der Waals forces and conventional hydrogen mostly, but there is an unfavorable donor-donor interaction observed in ARG19, which may explain its higher binding affinity value. Most notably, IL-6 (1ALU) has considerably less interactions compared to the other proteins. Ribociclib forms van der Waals contacts with THR140, GLN118, MET119 and VAL98, conventional hydrogen bonds with ASN146, and pi interactions with PRO141. Interestingly, ASN146 can form both conventional hydrogen bonds as well as partake in unfavorable donor-donor interactions, which can also explain the higher binding affinity value of IL-6 (1ALU). An unfavorable acceptor-acceptor interaction is observed in the Ribociclib-IL-17A (4HR9) complex, but the large number of Van der Waals interactions give a fairly negative binding affinity value. The interactions of Ribociclib with IFN-γ (1HIG) and IL-1β (9ILB) are mostly characterized by Van Der Waals forces, with the greater pi interactions in IFN-γ potentially attributing to lower binding affinity values. With CDK-6 (5L2T), Ribociclib interacts by forming conventional hydrogen bond with VAL101 and ASP163, amide-pi stacked with GLN103, pi-sigma with PHE98 and LEU152, pi-alkyl/alkyl with LYS43, VAL77, ALA41, ALA162, and VAL27. The dynamic stability of these interactions is investigated thoroughly through molecular dynamics simulations and estimation of binding free energy of interacting molecules.

### 3.3 Molecular Dynamics Simulation

MD trajectories of the apoproteins and protein-ligand complexes were analyzed for 100ns simulation time to check the changes in conformational structure of proteins. These provided detailed information on the dynamic stability of the proteins when forming protein-ligand complexes.

#### Root Mean Square Deviation (RMSD)

RMSD was used to observe the structural stability for nine complexes as well as their respective apoproteins. The root-mean-square deviation (RMSD) is significant as it quantifies the average distance between the atoms of superimposed protein structures, providing insight into the stability and conformational changes of the complexes over time. From Table 4, it can be observed that the average RMSD values of the complexes were favorably lower than 0.3nm (apart from the IFN-γ complex), indicating a stable conformation throughout the simulation period. Out of the eight experimental protein ligand complexes, five of the complexes with proteins MCP-1 -Ribociclib (0.1936±0.051), TNF-α -Ribociclib (0.2169±0.021), IL-1β -Ribociclib (0.1614± 0.026), IL-6 -Ribociclib (0.2447±0.026) and IL-17A/F -Ribociclib (0.2772±0.054) exhibited lower RMSD average values than that of their apoproteins. IFN-γ - Ribociclib (1.1583± 0.177), IL-17A -Ribociclib (0.2295 ± 0.032) and IL-17F -Ribociclib (0.3313 ± 0.062) complex showed higher RMSD value than its respective apoprotein. Notably, IL-17A/F (5N92) complex exhibited most reduced RMSD compared to its apoprotein, followed by MCP1 (1DOK) and IL-1β (9ILB). The RMSD of Ribociclib -CDK-6 complex was found to be slightly higher (0.2431 ± 0.036) than that of the apoprotein (0.2311 ± 0.030). From **Error! Reference source not found.**, we can observe fluctuations reach a stable during the course of the simulation. Additionally, it is observed that TNF-α (2AZ5), IL-6 (1ALU), IL-17A (4HR9), IL-17F (6HGO), and the control protein CDK-6 (5L2T) all reach stability after 60ns. This suggests that the ligand can form consistent and stable binding within the active sites of the proteins.

**Table 4.**
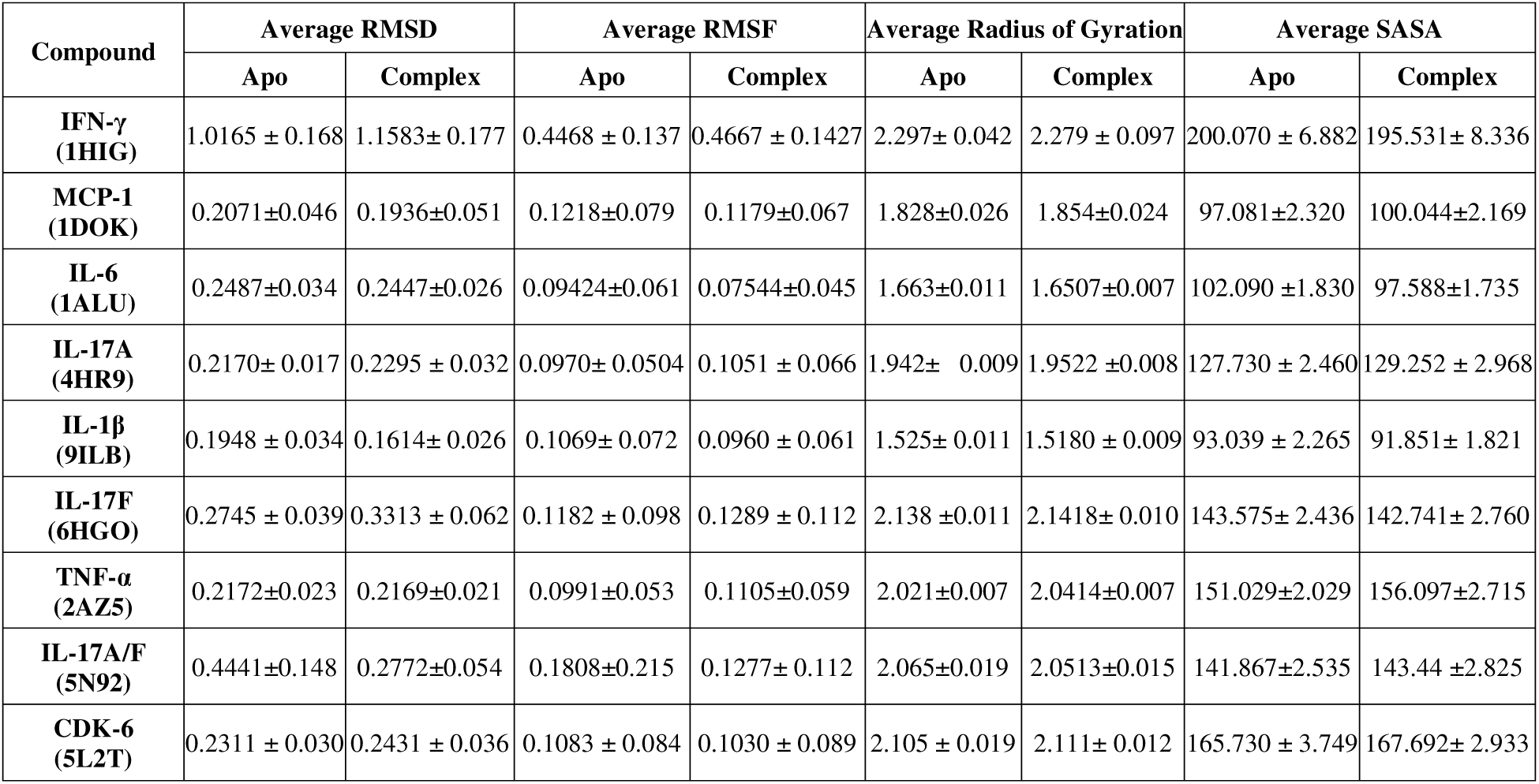
Average results of RMSD, RMSF, Rg and SASA of the Protein-ligand complex and their respective apoproteins.

#### Root Mean Square Fluctuation (RMSF)

The RMSF was calculated to determine the flexibility of individual amino acid residues in the protein. The root-mean-square fluctuation (RMSF) is crucial since it quantifies atomic fluctuations in a protein, offering insights into its flexibility and stability, while also revealing dynamic areas that may be essential for protein and ligand interactions. From Table 4, it can be noticed that the average RMSF value of eight protein-ligand complexes were less than 0.2nm. The complexes-IL-1β-Ribociclib (0.0960 ± 0.061), IL-6-Ribociclib (0.07544±0.045), IL-17A/F-Ribociclib (0.1277± 0.112), and MCP1-Ribociclib (0.1179±0.067) showed a decrease in average RMSF values compared to that of their native proteins as can be seen from Fig 4. However, the TNF-α -Ribociclib (0.1105±0.059), IL-17A-Ribociclib (0.1051 ± 0.066), IL-17F-Ribociclib (0.1289 ± 0.112), and IFN-γ -Ribociclib (0.4667 ± 0.1427) complexes showed greater average RMSF values than that of their native protein. The CDK-6 -Ribociclib (0.1030 ± 0.089) exhibited lower average RMSF value than that of apoprotein.

**Fig 4.**
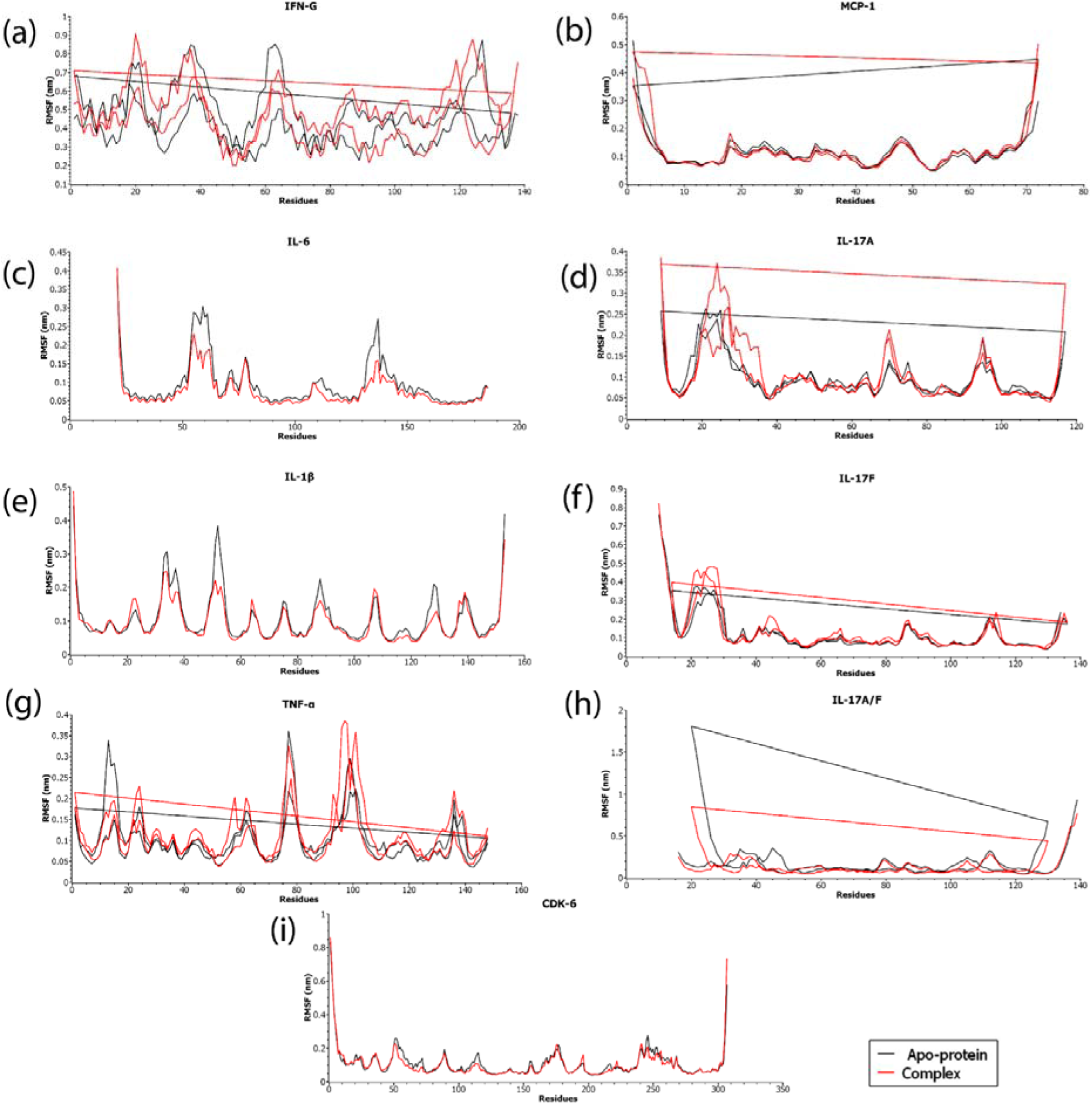
RMSF values plotted for apoprotein (black) and docked complexes (red) (a) IFN-γ (1HIG), (b) MCP-1(1DOK), (c) IL-6 (1ALU), (d) IL-17A (4HR9), (e) IL-1β (9ILB), (f) IL-17F (6HGO), (g) TNF-α (2AZ5), (h) IL-17A/F(5N92). and (i) CDK-6(5L2T).

**Fig 5.**
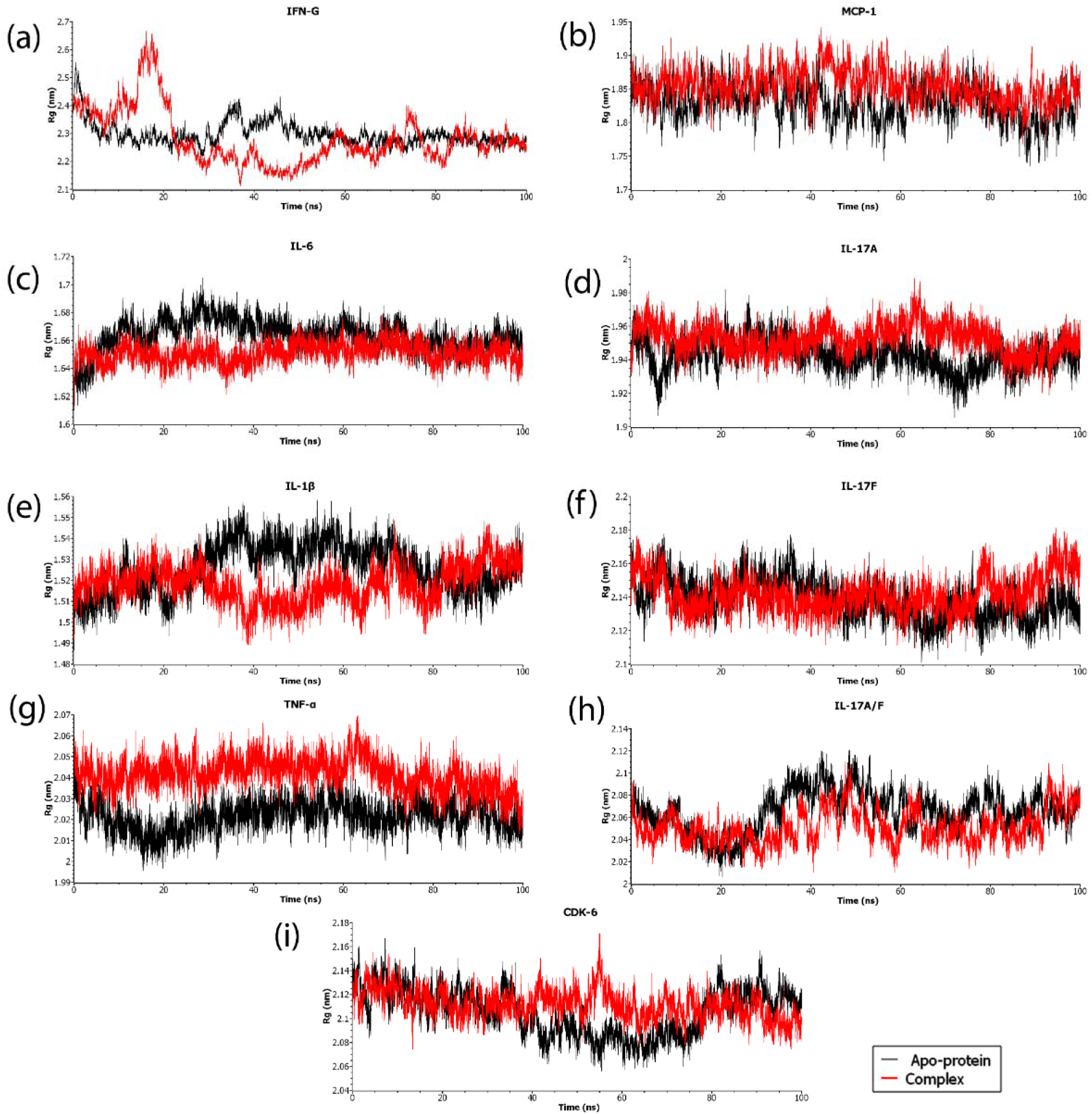
Rg values plotted for apoprotein (black) and docked complexes (red) (a) IFN-γ (1HIG), (b) MCP-1(1DOK), (c) IL-6 (1ALU), (d) IL-17A (4HR9), (e) IL-1β (9ILB), (f) IL-17F (6HGO), (g) TNF-α (2AZ5), (h) IL-17A/F(5N92). and (i) CDK-6(5L2T).

The individual RMSF values at the binding site were also evaluated and presented in **Error! Reference source not found.**1. It was found that regions corresponding to the active site and binding pocket exhibited lower RMSF values in general than that of their native proteins. In TNF-α, Ribociclib binding results in reduced RMSF values in the complex structure compared to the apo form for key interacting residues: CYS60 (apo 0.1102, complex 0.085), PRO104 (apo 0.105, complex 0.0924 and GLU107 (apo 0.0849, complex 0.1003). In IFN-G (1HIG), Ribociclib binding reduces the flexibility of several residues, indicating increased stability in the complex. ALA8, which participates in both conventional hydrogen bonding and Pi-sigma interaction, shows a slight increase in RMSF (apo 0.3806, complex 0.3986), while ARG131, involved in conventional hydrogen bonding, displays reduced flexibility (apo 0.3525, complex 0.3233). LYS12, forming a Pi-cation interaction, shows decreased RMSF (apo 0.4399, complex 0.396), and TYR53, involved in Pi–Pi stacking, also becomes more stable upon binding (apo 0.2831, complex 0.229). ALA17 (apo 0.4565, complex 0.402) and ILE73 (apo 0.2971, complex 0.3839), both engaged in Pi-alkyl interactions, show reduced and increased flexibility, respectively. Both IL-17A and IL-17F exhibited a higher average RMSF (Root Mean Square Fluctuation) in the Ribociclib-bound complex compared to their apo forms, indicating an overall increase in flexibility upon ligand binding. In IL-17F (6HGO), Ribociclib binding increases the flexibility of several residues such as VAL44 (apo 0.1226, complex 0.218) and PRO70 (apo 0.0781, complex 0.0947), that participate in Pi Alkyl/Alkyl bond..On the other hand, VAL74 involved in Pi Sigma bonding, displays reduced flexibility (apo 0.0773, complex 0.0769). TYR60, involved in Pi–Pi stacking, shows increased RMSF (apo 0.0788, complex 0.0826). VAL124 (apo 0.0639, complex 0.0557), VAL126 (apo 0.0573, complex 0.0544) and LYS121 (apo 0.063, complex 0.0617), involved in Van der Waals interactions, becomes more stable upon binding. GLN101 (apo 0.0732, complex 0.0795) and GLU102 (apo 0.0658, complex 0.0667), engaged in Conventional Hydrogen Bond and Carbon Hydrogen Bond/Pi-Donor Hydrogen Bond, show increased flexibility. Similarly in IL-17A (4HR9), LEU43 (apo 0.0808, complex 0.0968) and TRP57 (apo 0.0753, complex 0.0791) involved in Pi - Alkyl/Alkyl bonds show increased RMSF in the complex. Residues ASN26 (apo 0.1834, complex 0.3171), and GLN84 (apo 0.075, complex 0.0582), forms carbon hydrogen bonds with Ribociclib. SER30 (apo 0.1161, complex 0.1642), shows an unfavorable acceptor-acceptor interaction, while LYS104 (apo 0.0762, complex 0.0639), participates in a pi-cation interaction.

With CDK-6 (5L2T), Ribociclib interacts significantly through various residues. It forms conventional hydrogen bonds with VAL101 (apo 0.063, complex 0.0703) and ASP163 (apo 0.091, complex 0.061), and amide-pi stacked interactions with GLN103 (apo 0.0568, complex 0.0722). Additionally, Ribociclib establishes pi-sigma interactions with PHE98 (apo 0.0583, complex 0.0592) and LEU152 (apo 0.0467, complex 0.0526), as well as pi-alkyl/alkyl interactions with LYS43 (apo 0.0745, complex 0.0589), VAL77 (apo 0.0527, complex 0.0567), ALA41 (apo 0.0647, complex 0.0619), ALA162 (apo 0.0535, complex 0.0586), and VAL27 (apo 0.0847, complex 0.0734). The myriad interactions confirm overall rigidity and stability of these regions in the presence of Ribociclib.

#### Radius of Gyration (Rg)

The radius of gyration (Rg) was computed to describe the general compactness of the protein structures in both their apoprotein and complex forms. Comparisons between the values of Rg for each protein-ligand complex and the respective apoprotein states have shed light on the structural changes ensuing from ligand binding. From **Error! Reference source not found.**5, it can be observed that, in the complexes IFN-γ - Ribociclib (2.279 ± 0.097), IL-6 -Ribociclib (1.6507±0.007), IL-1β – Ribociclib (1.5180 ± 0.009), and IL-17A/F -Ribociclib (2.0513±0.015), the Rg is less than that of their native protein suggesting an increase in the protein’s compactness, indicating the stabilizing effect of Ribociclib. On the other hand, in complexes MCP1-Ribociclib (1.854±0.024), IL-17A – Ribociclib (1.9522 ±0.008), IL-17F -Ribociclib (2.1418 ± 0.010), and Tnf-α (0.2.0414 ± 0.007) exhibit slight increase in Rg which suggests a minor expansion in protein structure, though the protein remains relatively compact. Despite this, these complexes, like most other complex structures in this study, show stabilization within the 100 ns simulation period. A slight increase in the Rg can be observed for the complexes CDK-6 - Ribociclib (2.111± 0.012) and TNF-α - Ribociclib (2.0414±0.007) compared to the native protein. It also appears that the IFN-γ – complex structure destabilizes at the 60 ns mark but again tends to re-stabilize towards the end of the simulation. A much larger protein compared to other proteins in this study, the dynamic interactions of the IFN-γ (1HIG) may be better investigated with a longer simulation time.

#### Solvent-Accessible Surface Area (SASA)

The SASA was computed to assess the exposure of protein surfaces to solvent in both their apoprotein and ligand-bound forms. Higher SASA values indicate greater accessibility and flexibility of the structure to solvents. Conversely, lower SASA values imply reduced solvent exposure. For a protein-ligand complex, a lower SASA value compared to the apoprotein could potentially indicate the stabilizing effect of the ligand and increased protein-ligand interaction specificity.

Table 4 and Fig 6 compare the SASA values for each complex to their respective apo-protein states to give insights into the structural modifications that may arise when complying with ligand binding. IFN-γ – Ribociclib (195.531± 8.336), IL-6 -Ribociclib (97.588±1.735), IL-1β - Ribociclib (91.851± 1.821), and IL-17F -Ribociclib (142.741± 2.760), show a decrease in SASA upon binding to Ribociclib. The reduction in SASA values suggests that ligand binding reduces the exposure of the bounded protein to the solvent indicating a more compact structure. On the other hand, we can see a slight increase in SASA in these complexes of MCP1 – Ribociclib (100.044±2.169), IL-17A – Ribociclib (129.252 ± 2.968), IL-17A/F -Ribociclib (143.44 ±2.825), CDK-6 -Ribociclib (167.692± 2.933) and TNF-α -Ribociclib (156.097±2.715), indicating a minor expansion in the protein’s surface area upon binding with Ribociclib.

**Fig 6.**
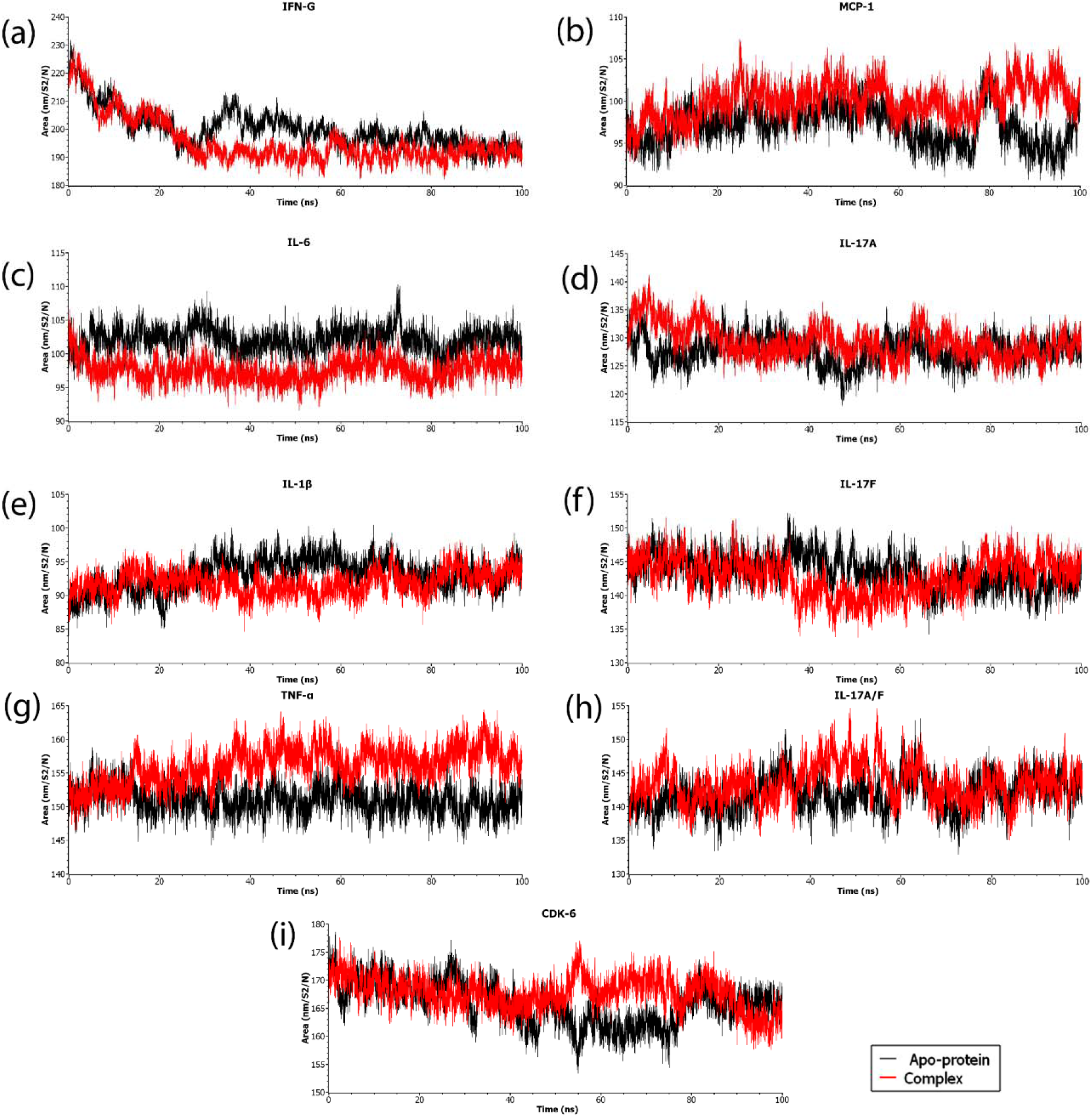
SASA values plotted for apoprotein (black) and docked complexes (red) (a) IFN-γ (1HIG), (b) MCP-1(1DOK), (c) IL-6 (1ALU), (d) IL-17A (4HR9), (e) IL-1β (9ILB), (f) IL-17F (6HGO), (g) TNF-α (2AZ5), (h) IL-17A/F(5N92). and (i) CDK-6(5L2T).

**Fig 7.**
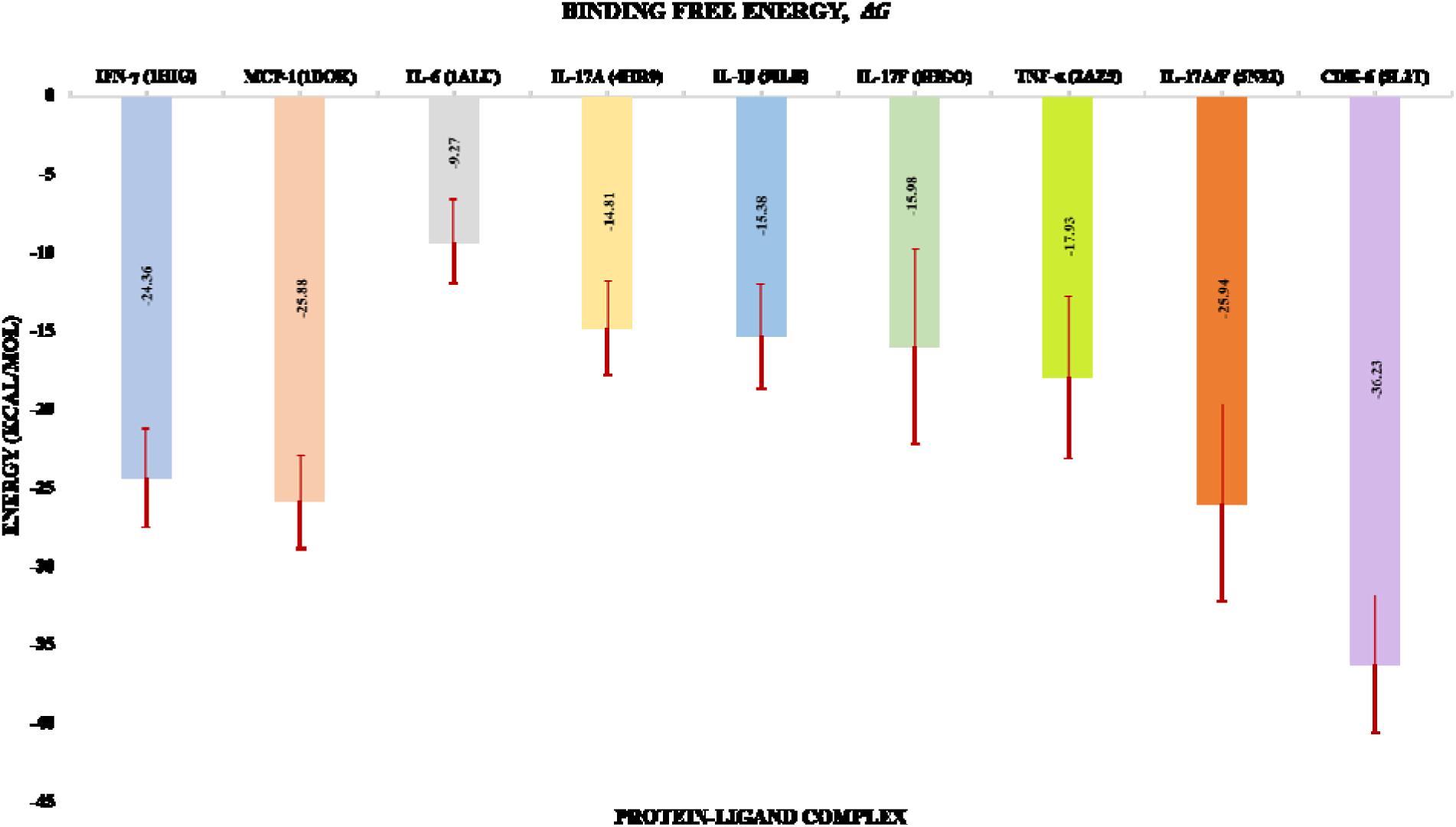
Total binding free energy of the complexes

**Fig 8.**
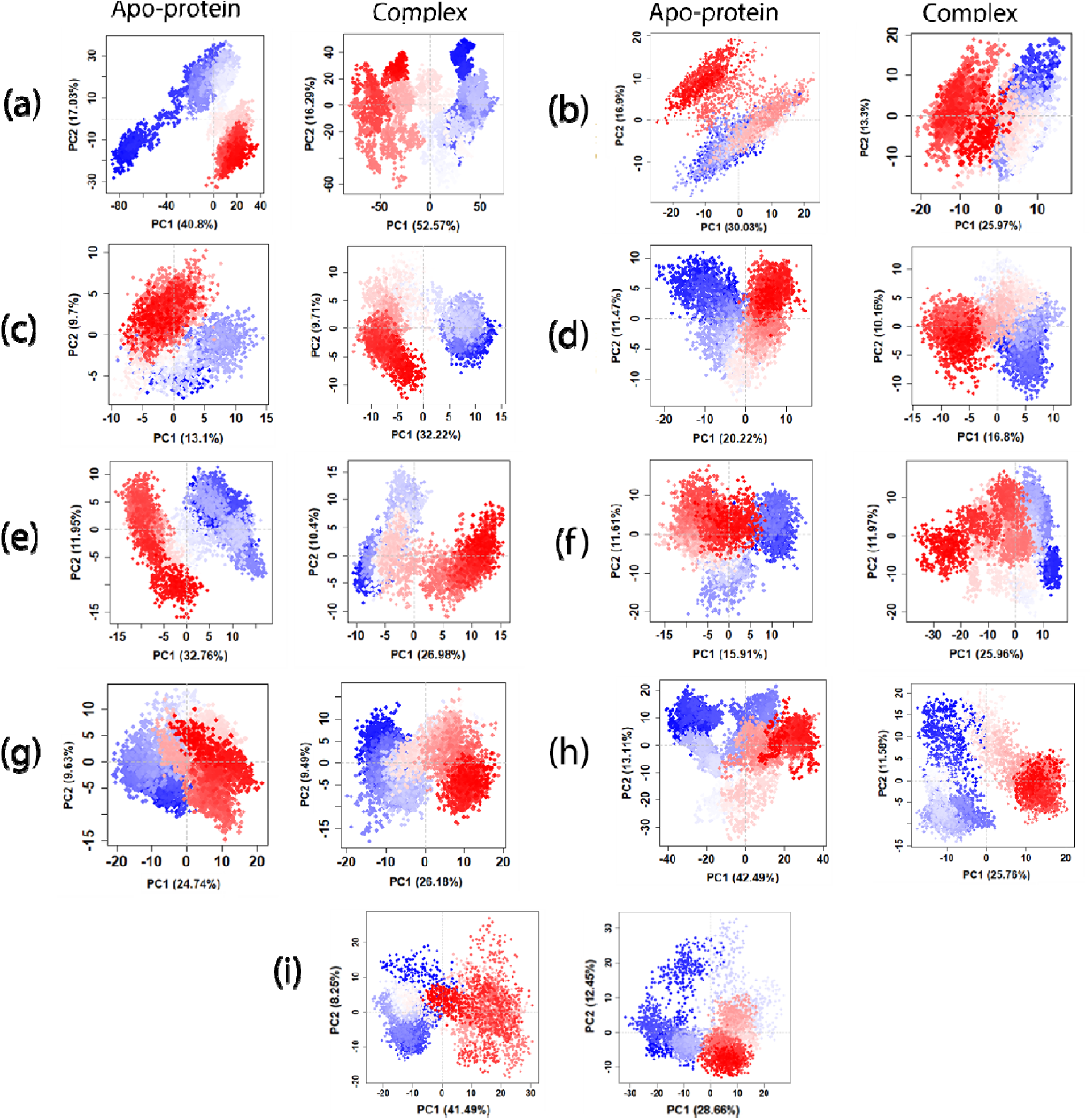
Principal component analysis of the MD simulation trajectories for apoprotein and docked complexes (a) IFN-γ (1HIG), (b) MCP-1(1DOK), (c) IL-6 (1ALU), (d) IL-17A (4HR9), (e) IL-1β (9ILB), (f) IL-17F (6HGO), (g) TNF-α (2AZ5), (h) IL-17A/F(5N92). and (i) CDK-6(5L2T).

**Fig 9.**
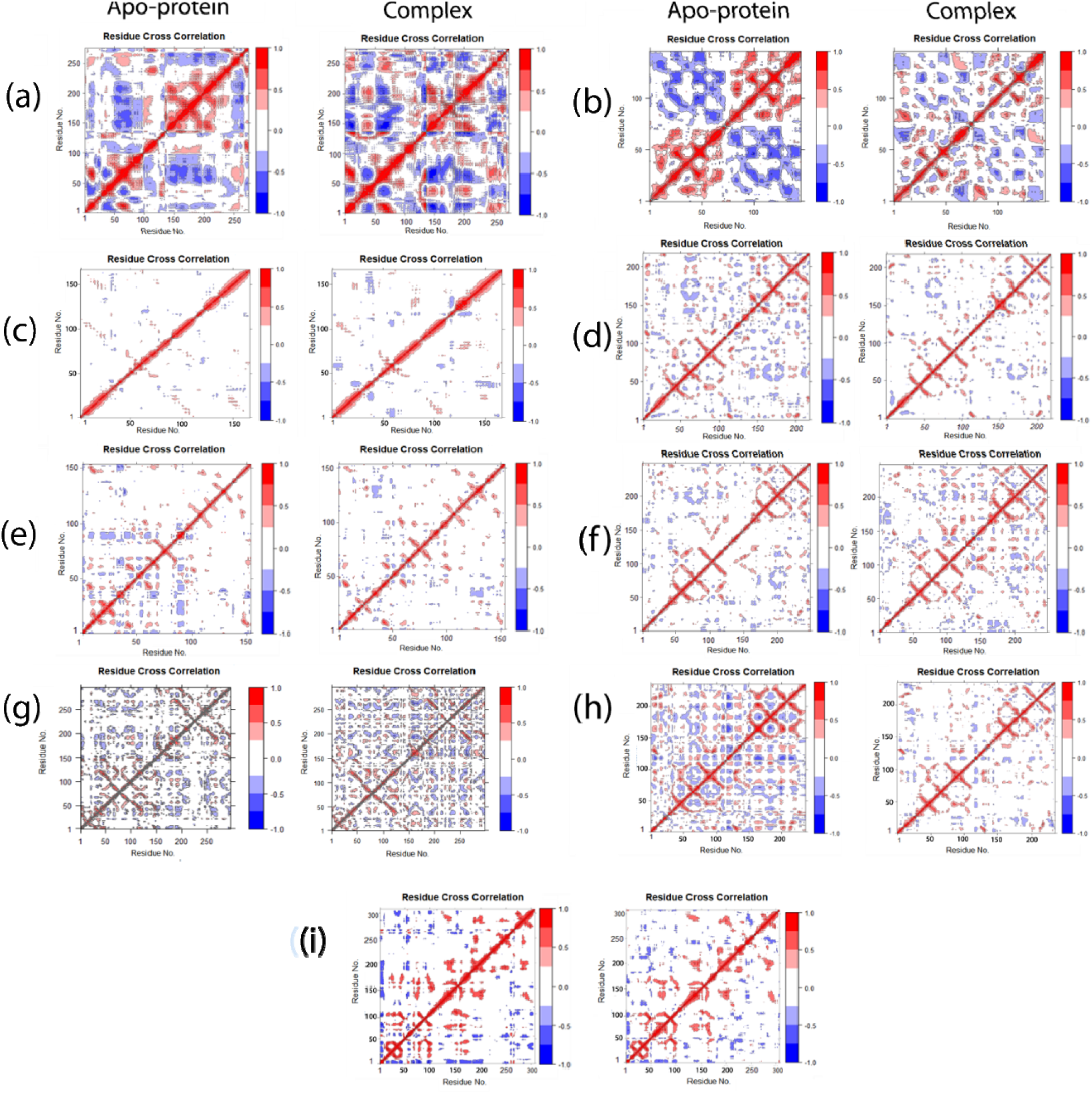
DCCM analysis for apoprotein and docked complexes (a) IFN-γ (1HIG), (b) MCP-1(1DOK), (c) IL-6 (1ALU), (d) IL-17A (4HR9), (e) IL-1β (9ILB), (f) IL-17F (6HGO), (g) TNF-α (2AZ5), (h) IL-17A/F(5N92). and (i) CDK-6(5L2T).

### 3.4 MMGBSA

The MMGBSA method was applied to investigate the binding free energies of the cytokines in complex with their respective ligands. The last 50 ns of the trajectory was analyzed with binding energy values rangin from −9.27 kcal/mol to −36.23 kcal/mol. From the results in **Error! Reference source not found**.7 and Table 5, we can see that the pro-inflammatory cytokines IL-17A/F (5N92) and MCP-1 (1DOK) show the strongest binding affinities apart from the control protein, while IL-6 (1ALU) shows the weakest. IL-17A/F’s overall binding energy is −25.95 kcal/mol, which is dominated by Van der Waals and electrostatic interactions. The electrostatic interactions stem from the presence of conventional and pi-donor hydrogen bonds and is second only to those found in Ribociclib-IL-17F complex. MCP-1 has a total binding energy of −25.85 kcal/mol, with moderate contributions from both Van der Waals forces and electrostatic interactions. However, the comparatively lower overall solvation energy (third overall after IL-6 and IFN-γ) gives this interaction sufficiently low total binding energy. IL-6 (1ALU) has a total binding energy of −9.27 kcal/mol, with the lowest contribution from Van der Waals forces compared to the other complexes. The largest Van Der Waals contribution is seen in the Ribociclib-IFN-γ (1HIG) complex, but the complex has the weakest electrostatic interactions among all proteins studied. IL-17F shows the strongest electrostatic interactions due to the presence of hydrogen bonds, pi-sigma and pi-pi stacked interactions. However, the complex exhibits moderate Van Der Waals interactions and high solvation energy and has a total binding free energy of −14.81 kcal/mol. TNF-α, on the other hand, show relatively stronger Van Der Waals interactions but moderate electrostatic interactions and solvation energy, yielding a total binding energy of −17.93 kcal/mol. IL-1β and IL-17F exhibit moderate interactions with total binding energies of −15.38 kcal/mol and −15.98 kcal/mol, respectively. The control protein, CD6-K (5L2T) exhibited strongest interactions with the lowest total binding free energy of −36.23 kcal/mol dominated by the strongest Van Der Waals interactions, strong electrostatic interactions, and low solvation energies, confirming Ribociclib’s role as a CDK-6 inhibitor. The raw data of binding affinity calculations are summarized in Table S2.

**Table 5.**
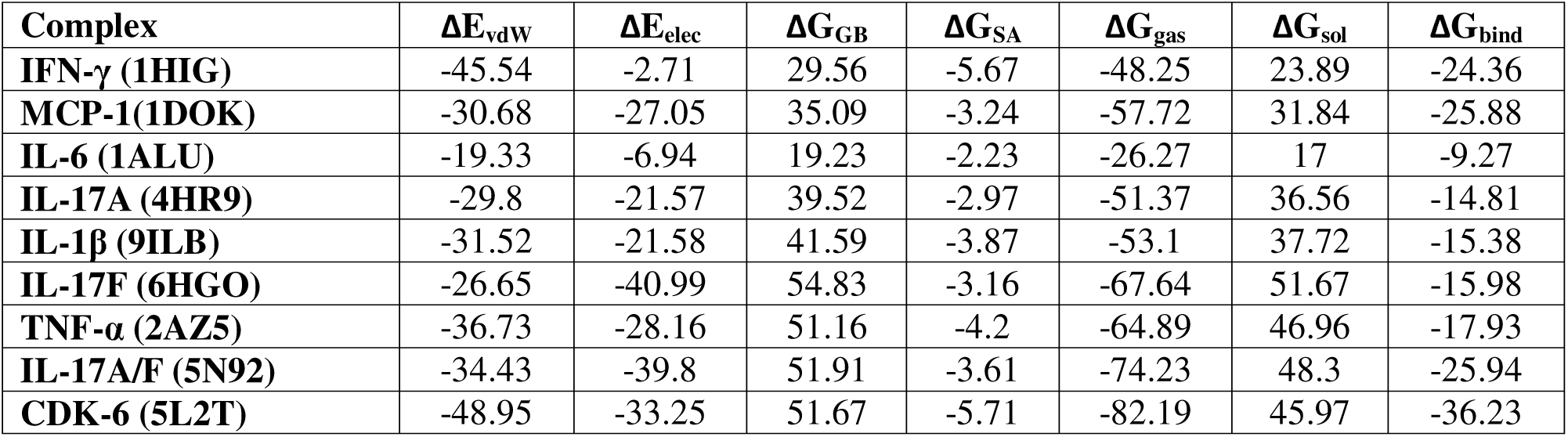
The calculation of binding free energy results of Ribociclib in 8 protein complexes (all energy terms are in kcal/mol)

### 3.5 Principal component analysis (PCA) & Dynamic cross-correlation matrix (DCCM)

Principal component analysis (PCA) was applied to the last 50 ns of the MD trajectories for apo forms of the proteins and the protein-ligand complexes in order to shed light on the conformational dynamics and structural differentiation. **Error! Reference source not found.**8 depicts how the conformational states of the proteins and the protein-ligand complexes cluster with respect to the top two principal components (PCs). Each dot represents a step in the simulation with blue denoting initial, unstable conformations, white denoting intermediate time steps, and red denoting stable, later conformations. Overall stability of a structure is determined by the size and spread of the cluster, with lower spread and higher clustering of a particular color being associated with overall stable structure during the simulation period. It can be observed that the protein-Ribociclib complexes of IL-17A/F, IL-6, MCP-1, TNF-α and CDK-6 show dense red clustering along a particular PC. The complex of IFN-γ, despite having similar binding free energies to IL-17A/F and MCP-1, give a comparatively wider spread of conformational values, suggesting flexibility in the protein backbone despite the stabilizing effects of Van Der Waals interactions in the presence of the ligand. This observation may be attributed to the differences in size as IL-6, the smallest protein in the cohort, exhibit conformational stability despite the highest binding free energy values. Apart from IFN-γ and IL-6, the results are generally in line with our observation from the MM-GBSA experiment where narrow, dense clustering is associated with lower binding free energy values.

In **Error! Reference source not found.**9, the DCCM graphs show the correlation coefficients between residue pairs for each protein in both apo and complex forms. Positive correlations (red) indicate cohesive movements, while negative correlations (blue) suggest anti-correlated movements. The complexes MCP-1 –Ribociclib, IL-17A/F –Ribociclib, and CDK-6 – Ribociclib (control group) exhibit stronger positive correlations compared to their apo forms, indicating a general trend of structural stabilization upon ligand binding. Compared to their apo structure, the complexes of IFN-γ, IL-6, IL-17F, and TNF-α showed an increase in both positive and negative correlations indicating significant structural changes due to the formation of the complex. In contrast, the complexes of IL-17A and IL-1β did not exhibit any marked increase in the proportion of positive correlations compared to their apoprotein forms. These results are also consistent with the findings and evaluations from MM/GBSA and PCA.

This work provides a comprehensive assessment of the inhibitory activities of the small molecule Ribociclib against the major classes of pro-inflammatory cytokines. The inclusion of a control protein and known receptor of Ribociclib, CDK-6, enabled us to identify the most important metrics to evaluate the inhibitory performance of Ribociclib. Binding free energy estimations, principal components analysis, and dynamic cross-correlation matrix emerged as the best estimators of Ribociclib activity based on its performance against the CDK-6 protein, which gave the best results for all three metrics. CDK-6 was followed by IL-A/F and MCP-1, which had the second and third lowest binding free energy values (after CDK-6), respectively and demonstrated conformational stability in PCA and DCCM analysis. Ribociclib exhibited moderate activity against TNF-α, IL-6, and IFN-γ. TNF-α ranked fifth in terms of binding free energy estimates but had good-to-moderate results in the PCA and DCCM analysis. IL-6 had the lowest binding free energy values but demonstrated good conformational stability in PCA and moderate results in DCCM analysis. Ribociclib-IFN-γ complex was stabilized by strong Van Der Waals interactions but did not show marked performance in PCA and DCCM analysis. Ribociclib demonstrated the least inhibitory activities against IL-17F, IL-1β, and IL-17A. Interestingly, the Ribociclib complex of IL-17F exhibited the strongest electrostatic interactions which prompted us to include the heterodimer IL-17A/F into our analysis, which was not included in the preliminary evaluations. Despite the relatively low inhibitory activities against the homodimers IL-17A and IL-17F, Ribociclib showed strong inhibitory activities against the heterodimer IL-17A/F. Experimental studies are required to confirm our analysis if this inhibition is strong enough to prevent receptor binding of IL-17A/F and result in the subsequent arrest of inflammatory pathways implicated in autoimmune and infectious diseases (55).

Ribociclib exhibits a broad range of molecular interactions against the target proteins. and pyridopyrimidine core structure increases the ability of Ribociclib to form these interactions due to their aromaticity and electron-rich nature, which remarkably increases the stability and specificity of the binding of Ribociclib, making it a potential inhibitor of multiple pro-inflammatory cytokines. This is a structural characteristic which is similar to other kinase inhibitors, such as Ruxolitinib and Tofacitinib, which also depend on their distinctive aromatic and cyclic structures for making key interactions with their targets. For example, Ruxolitinib, a pyrazole containing a 2-cyano-1-cyclopentylethyl group and a pyrrolo[2,3-d]pyrimidin-4-yl group, and Tofacitinib, a pyrrolopyrimidine bearing an N-methyl,N-(1-cyanoacetyl-4-methylpiperidin-3-yl) amino moiety, use their aromatic rings and electron-rich groups to interact with non-specific protein-tyrosine kinases, in a similar way that Ribociclib interacts with cytokines. It is important to note that the estimated binding affinities of Ribociclib with the target proteins may not be directly comparable to those of ligands specifically designed to target these individual proteins (56,57). However, in complex inflammatory responses involving multiple cytokines, such as those observed in autoimmune disorders or viral infections, the use of a versatile inhibitor like Ribociclib may offer a therapeutic advantage over conventional single-target agents. The findings of our work suggest several promising Ribociclib targets and provide a foundation for further experimental validation and possible clinical applications.

## 4. Conclusion

In this work, Ribociclib’s potential as a common inhibitor of eight significant classes of pro-inflammatory cytokines, emphasizing its ability to reduce inflammatory responses, was investigated. In silico methods such as ADMET analysis, molecular docking, molecular dynamics simulations, MMGBSA, PCA, and DCCM analysis were conducted to assess the interactions between Ribociclib and cytokines. Molecule database was thoroughly examined for drug-like characteristics and conducted molecular docking experiments to discover favorable binding interactions. The stability of these interactions was validated by further validation using RMSD, RMSF, Rg, and SASA analyses in MD simulations. The DCCM and PCA analysis revealed further information on the protein-ligand complexes’ dynamic behavior. Notably, Ribociclib exhibited substantial binding to IL-17A/F, and MCP-1 proteins, suggesting favorable affinities and improved structural stability. The small molecule also exhibited moderate binding with TNF-α, IL-6, and IFN-γ. Findings from this research indicate that Ribociclib may be a promising inhibitor of pro-inflammatory cytokines, providing a possible treatment method for inflammatory illnesses. The aromatic benzene rings of Ribociclib, which offer π interactions, share structural features with other kinase inhibitors, such as Ruxolitinib and Tofacitinib, hence underlying its ability to interact with multiple classes of cytokines. Additional research, such as extended molecular dynamics simulations and collaboration with experimental researchers for in vitro and in vivo studies, is required to confirm these outcomes and transform them into beneficial outcomes.

## Competing Interests

The authors declare no competing interests.

## Funding Information

This research did not receive any specific grant from funding agencies in the public, commercial, or not-for-profit sectors.

## Author Contributions

Akid Ornob conceptualized and designed the study. Rabita Rahman Era acquired the data and performed data analysis. Both authors contributed to the interpretation of the data, and to the drafting and revision of the final manuscript. Both authors read and approved the final version of the article prior to submission.

## Data Statement

All data supporting the findings of this study are included within the main article and the supplementary information files.

## Acknowledgements

The authors express their gratitude to the Department of Biomedical Engineering at the Military Institute of Science and Technology (MIST) for providing access to its computational facilities. The authors also express their appreciation to Israt Sultana and Saira Binte Awal for their valuable assistance during the preliminary phase of this investigation.

## Notes

### Competing Interest Statement

The authors have declared no competing interest.

