## Supplementary Figures and Supplementary Tables S1 and S2 for "Ribociclib as a Potential Multi-Target Inhibitor of Pro-Inflammatory Cytokines: An In Silico Investigation": Supplementary Figures.docx

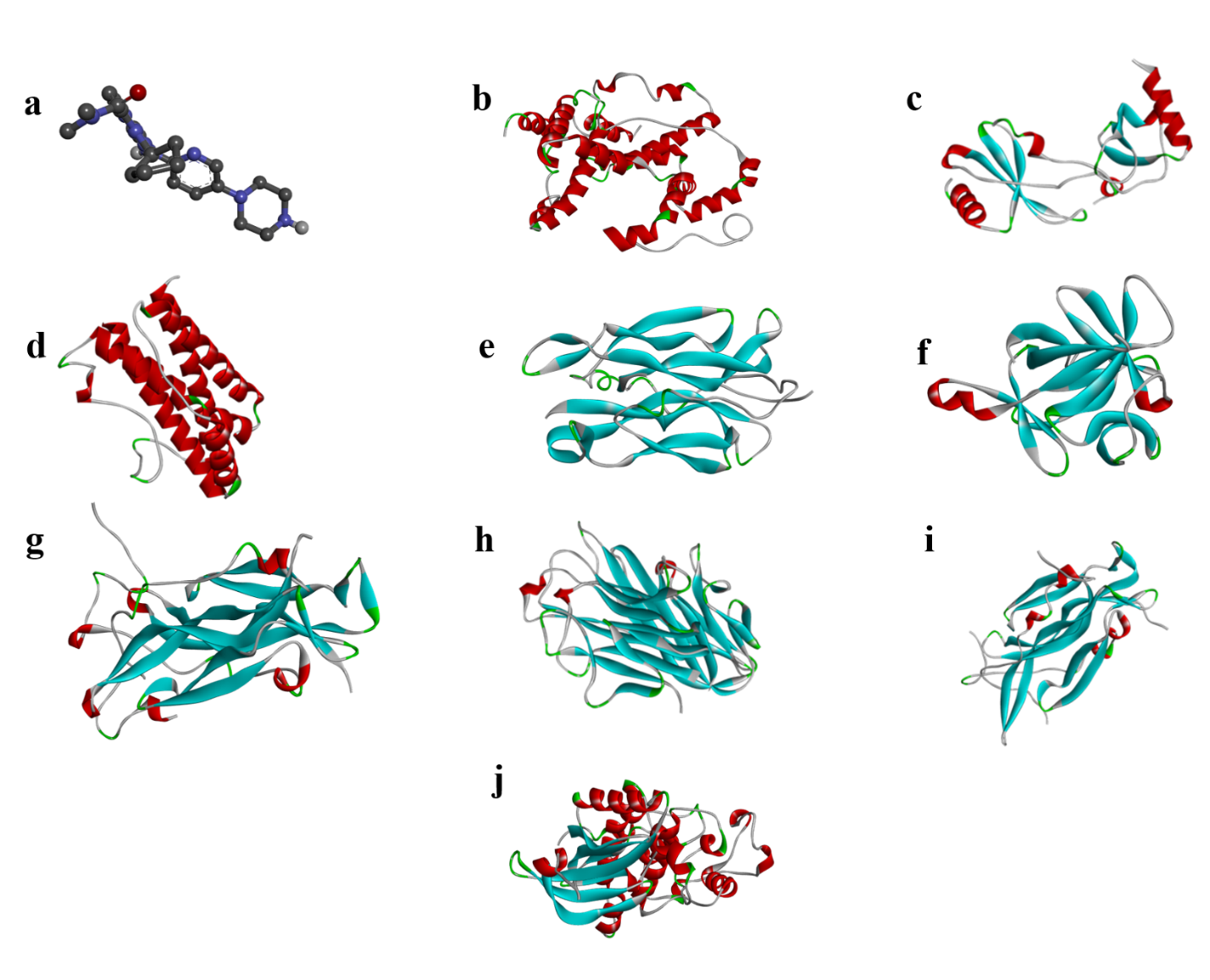
Fig S1. Structures of (a)Ribociclib, (b) IFN-γ (1HIG), (c) MCP-1(1DOK), (d) IL-6 (1ALU), (e) IL-17A (4HR9), (f) IL-1β (9ILB), (g) IL-17F (6HGO), (h) TNF-α (2AZ5), (i) IL-17A/F(5N92). and (j) CDK-6(5L2T).

Fig S2. Molecular docking of Ribociclib with 7 targeted proteins. (a) IFN-γ (1HIG), (b) MCP-1(1DOK), (c) IL-6 (1ALU), (d) IL-17A (4HR9), (e) IL-1β (9ILB), (f) IL-17F (6HGO), (g) TNF-α (2AZ5), (h) IL-17A/F(5N92). and (i) CDK-6(5L2T). Interacting residues of Ribociclib are shown by ball and sticks colored by elements

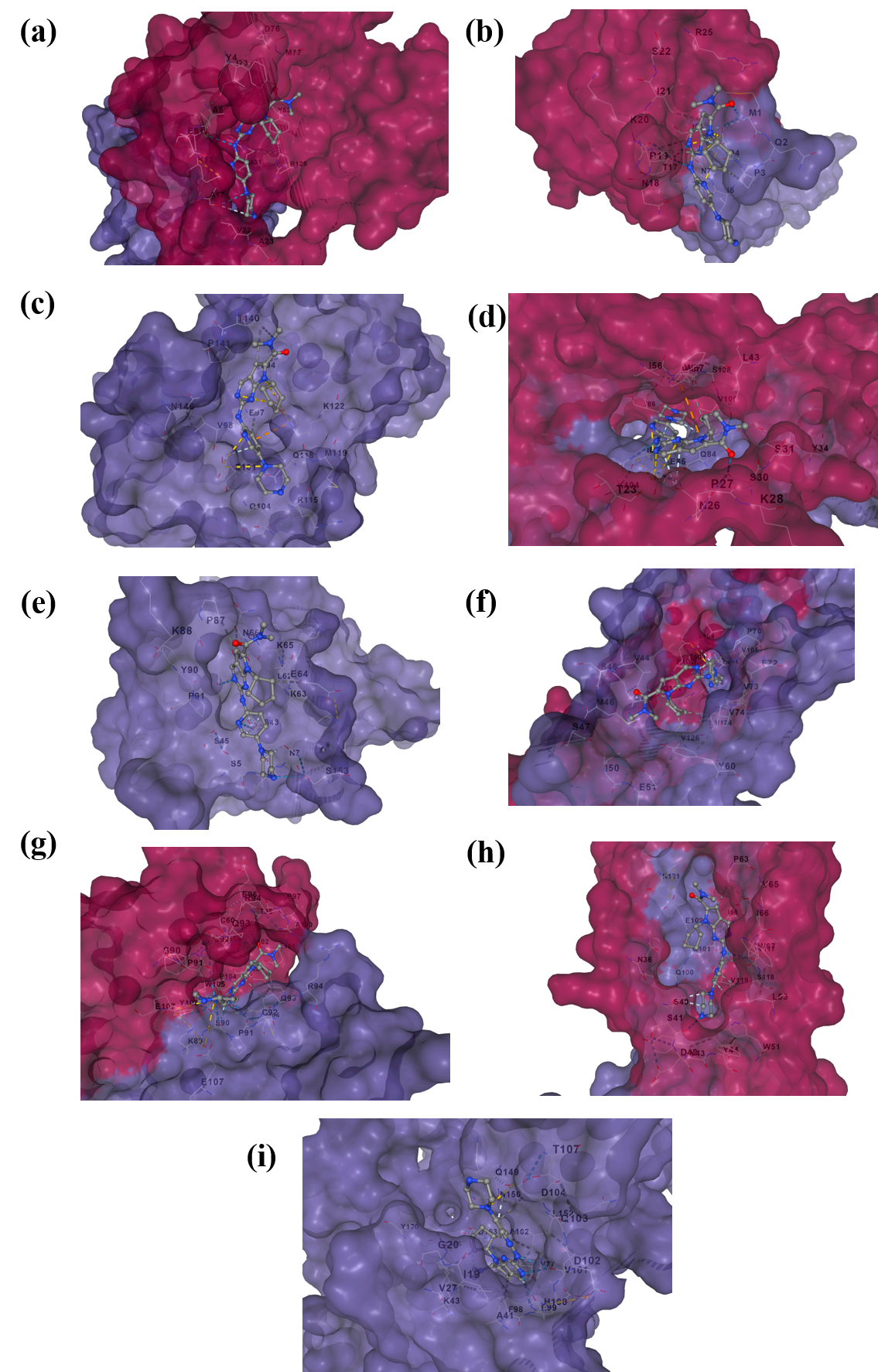
